# Paperfuge: An ultra-low cost, hand-powered centrifuge inspired by the mechanics of a whirligig toy

**DOI:** 10.1101/072207

**Authors:** M. Saad Bhamla, Brandon Benson, Chew Chai, Georgios Katsikis, Aanchal Johri, Manu Prakash

## Abstract

Sample preparation, including separation of plasma from whole blood or isolation of parasites, is an unmet challenge in many point of care (POC) diagnostics and requires centrifugation as the first key step. From the context of global health applications, commercial centrifuges are expensive, bulky and electricity-powered, leading to a critical bottle-neck in the development of decentralized, electricity-free POC diagnostic devices. By uncovering the fundamental mechanics of an ancient whirligig toy (3300 B.C.E), we design an ultra-low cost (20 cents), light-weight (2 g), human-powered centrifuge that is made out of paper (“paperfuge”). To push the operating limits of this unconventional centrifuge, we present an experimentally-validated theoretical model that describes the paperfuge as a non-linear, non-conservative oscillator system. We use this model to inform our design process, achieving speeds of 125,000 rpm and equivalent centrifugal forces of 30,000 g, with theoretical limits predicting one million rpm. We harness these speeds to separate pure plasma in less than 1.5 minutes and isolate malaria parasites in 15 minutes from whole human blood. By expanding the materials used, we implement centrifugal microfluidics using PDMS, plastic and 3D-printed devices, ultimately opening up new opportunities for electricity-free POC diagnostics, especially in resource-poor settings.

## Introduction

A centrifuge is the workhorse of any medical diagnostics facility. From extraction of plasma from whole blood (for performing immunoassays or determining hematocrit value), to concentration of pathogens and parasites in biological fluids such as blood, urine and stool (for microscopy), centrifugation is the first key-step for most diagnostic assays (1). In modern diagnostics, separation of unwanted cellular debris is especially critical for the accuracy and reliability of molecular diagnostics tools and lateral flow-based rapid diagnostic tests (RDT) (2) that are designed for detecting low-levels of infections in diseases such as malaria, HIV and tuberculosis (3–5). Currently, centrifugation is inaccessible in field conditions since conventional machines are bulky, expensive and require electricity (4). The need for electricity-free centrifugal bio-separation devices prompted researchers to use egg-beaters and salad-spinners as proposed devices (6,7). However, these suffer from bulky design and extremely low rotational speeds (max 1,200 rpm; 300 g), leading to impractical centrifugation times for a simple task of blood plasma separation (*>* 10 mins). Thus, a low-cost, portable, human-powered centrifuge that achieves high-speeds is an essential, yet unmet need, especially for diagnostics in resource-limited environments (8–10).

Here we describe the design and implementation of an ultra-low cost (*<* 20 cents), light-weight (2 g), field-portable centrifuge, henceforth referred to as *paperfuge*. We demonstrate that the paperfuge achieves speeds of 125, 000 rpm (30, 000 g) using only human power. Using a combination of modeling and experimental validation, we uncover the detailed mechanics of the paperfuge and leverage this understanding to construct centrifuges from different materials (paper, plastic etc.). We demonstrate applications including plasma separation, quantitative buffy coat analysis (QBC) and integrated centrifugal microfluidic devices for point-of-care (POC) diagnostic testing.

## Results and Discussion

We start by measuring the speed of a buzzer made out of a paper-disc or paperfuge (Fig. 1B, *R*_d_ = 50 mm), using a high-speed camera (*f_rec_* = 6, 000 fps, Fig. 1C, Fig. S3, movie S1 and S2). The actuation of the paperfuge consists of successive “unwinding” and “winding” phases (Fig. 1D). In the unwinding phase, the outward input force (applied by human hands on the handles) accelerates the disc to a maximum speed 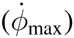. In the winding phase, the input force falls to zero, allowing the disc’s inertia to rewind the strings and draw the hands back inwards. Since the strings are flexible (low bending stiffness), they wind beyond the geometric zero-twist point (58), passing through a spectrum of helical twisting states. After reaching a tightly-packed super-coiled state (13, 14) (Fig. 1C inset), the motion of the disc comes to a momentary halt. At this point, an outward force is re-applied, unwinding and winding the strings. This cycle repeats itself at a frequency (*f*_0_). Fig.1 E shows the rotational speeds 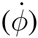 of paperfuges sizes ranging over an order of magnitude (5 to 85 mm).

**Figure 1.**
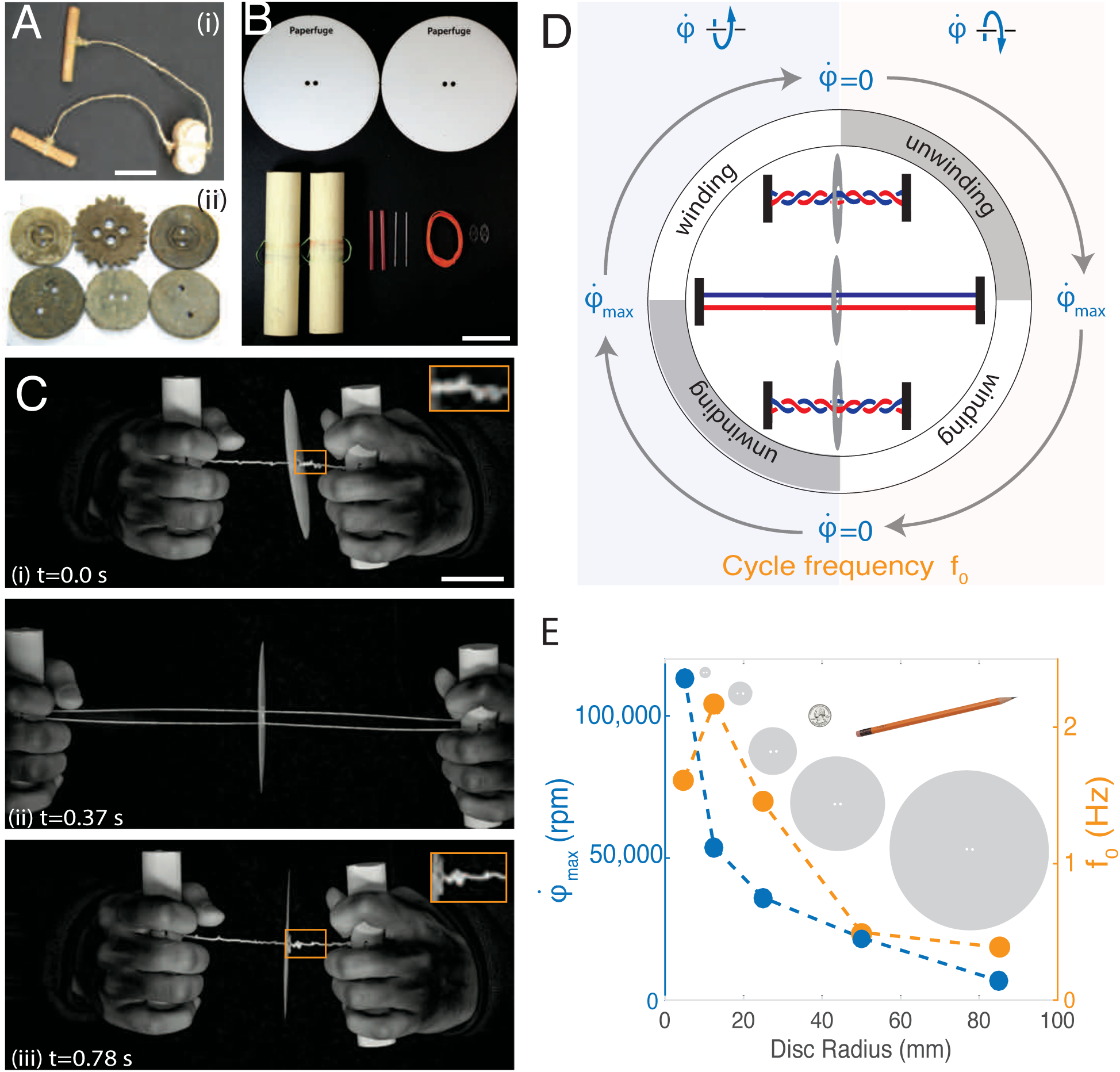
Spinning dynamics of a paperfuge. **A)** (i) Images of an ancient Caribou bone tied to a raw-hide, used as a charm (40), and (ii) lead whirligigs relics with saw-tooth edges (36). **B)** Materials used to construct paperfuge include paper discs, wooden handles, string, capillaries, capillary holders and plastic shims (see Methods for details). **C)** High-speed images of a rotating paperfuge showing a succession of (i) wound, (ii) unwound, and (iii) a final wound state, for half a cycle. The entire process occurs in a second (movie S1). The insets show the supercoiling of the strings during the wound states. **D)** Schematic illustrating the periodic rotation of the paperfuge where the disc (shown in gray) starts at zero rotational speed 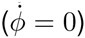 and the cycles to maximum speed 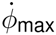. **E)** Plot of 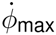 and *f*_0_ for different human-powered discs (*R_d_* = [5, 12.5, 25, 50, 85] mm, Table S3). We find that the radius of the central disc (*R*_d_) strongly influences the maximum speed 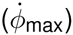 achieved by the disc. The smallest disc (*R*_d_ = 5 mm) reach 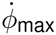 = 125, 000 rpm at a *f*_0_ = 2.2 Hz (Fig. 1E and). It is worthwhile noting muscle force-velocity constrain the fastest operable frequency at 2 Hz (11). To the best of our knowledge, 125, 000 rpm is the fastest rotational speed reported via a human-powered device (12). The background shows images of disc sizes along with a U.S. quarter and a pencil for visual comparison. Scale bars are 5 cm.

By optimizing the buzzer for high-speed operation and driving it using human hands, we achieve roughly two orders of magnitude faster rotation speeds than previously reported (15). Motivated by this experimental finding, we develop a detailed theoretical model that faithfully captures the extensive parameter-space of this system.

We describe the buzzer toy as a non-linear, non-conservative oscillator: in every rotation cycle, the input energy is introduced by human hands (applied force) and is dissipated by the system through air drag and in the strings. We consider a massless, inextensible string with uniform tension throughout the string (Fig. 2A). Starting from an energy balance on the system (see SI), we derive a governing equation of motion that relates angular acceleration 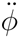 and different torque components as,

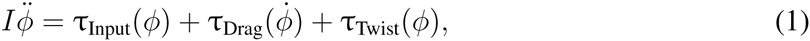

where *I* is the inertial moment of the disc, *ϕ* is the angular displacement, and τ_Input_, τ_Drag_, τ_Twist_ are the input, air-drag and string-twisting contributions to the torque, respectively. Each of the above terms can be derived from the geometrical parameters of the system (see SI for details).

**Figure 2.**
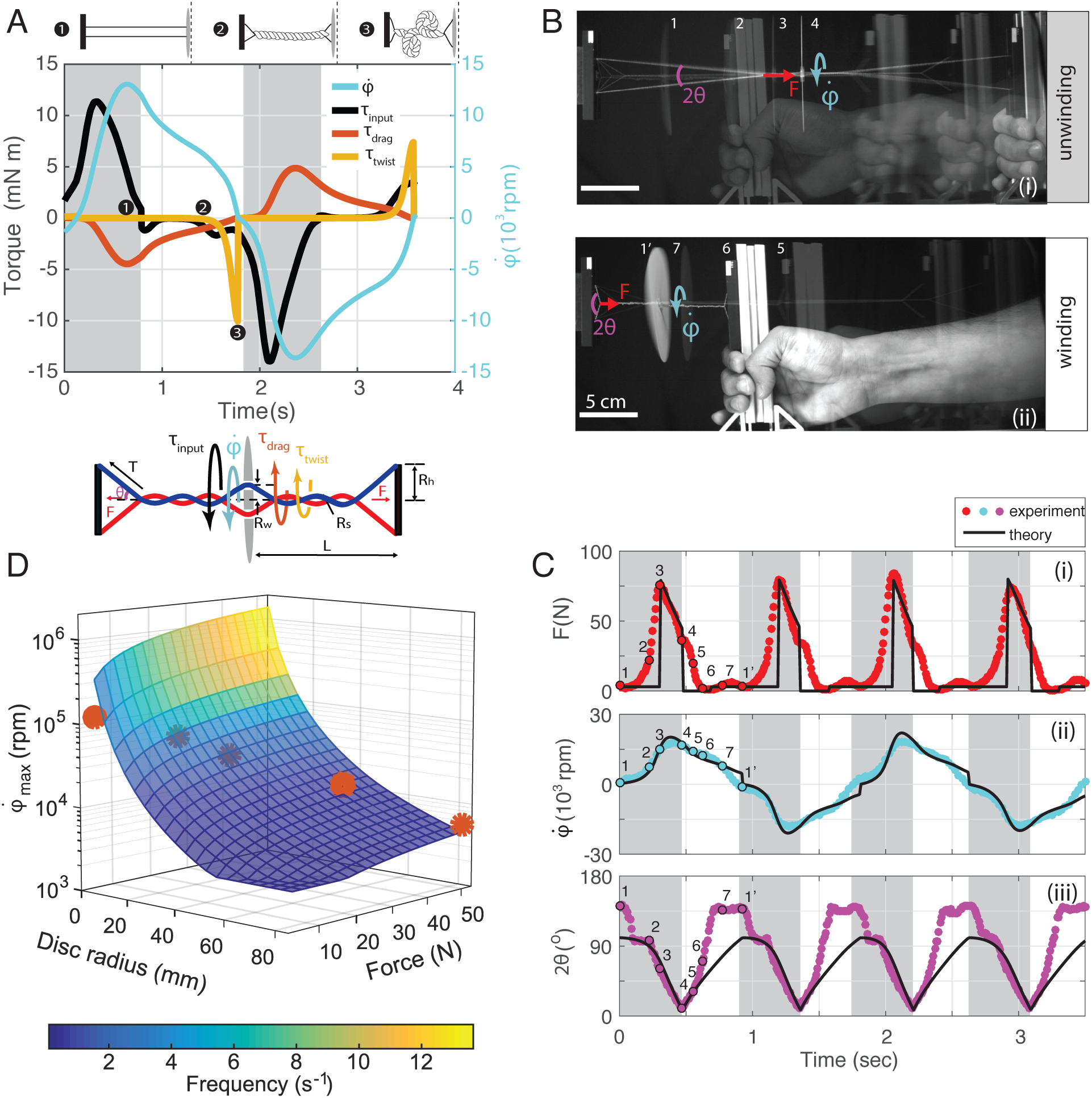
Validation of theoretical model with experiments. **A)** Numerical solution (see SI section for details) highlighting contributions of different torque components. Overlayed schematics illustrate the coiling-states and the geometry of the paperfuge. **B)** (i) Image of four (1 – 4) superimposed snapshots of the paperfuge during the “unwinding phase” where a force *F* is applied, unwinding the coiled-strings and accelerating the paperfuge to its maximum rotational speed (SI methods and movie S4). (ii) Four (5 – 1′) superimposed image showing the “winding phase” where the force *F* falls to zero, and the inertia of the disc causes re-winding of the strings. **C)** Synchronized time series plots of (i) force *F*, (ii) the rotational speed 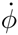 of the disc, and (iii) the angle 2*θ* of the strings. The colored circles denote experimental data and the continuous black lines denote the predictions from theory, which are in good agreement. When *F* ∼ 0, low string tension causes the strings to slacken, assuming a geometry outside the scope of our model, causing a systematic underprediction in *θ*. **D)** Choosing equivalent design parameters, we show a theoretical design landscape with overlayed experimental data-points (*R_d_* = [5, 12.5, 25, 50, 85] mm, red circles). The scaling curve predicts that a million rpm can be achieved at a frequency of oscillation of *>* 10 Hz and nominal forces (50 N).

We first start with the input torque that is dependent on the applied pull force as τ_Input_(*ϕ*) = −sgn(*ϕ*)2*R_s_F* tan *θ* where *R_s_* is the radius of the string, and *θ* is the angle subtended by the string from the axial axis of the paper disc. To evaluate τ_Input_(*ϕ*), we must find a relation between *θ* and *ϕ*. We divide the twisting section of the string into three parts (*l*_1_, *l*_2_, *l*_3_, Figs. S4 and S5), and obtain a geometrical relationship between *θ* and *ϕ*, where *θ* = sin^−1^(|*ϕ*|*R_s_* + *R_h_* + *R_w_*)/*L*), *R_h_* is the handle radius, 2*R_w_* is the distance between the holes on the disc, and 4*L* is the total length of the string. This gives the formulation 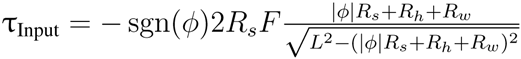.

Since the Reynolds number for the spinning disc is large (*Re* ∼ 10^5^), we calculate the component due to air-drag as 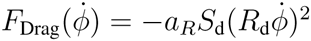, where *a_R_* is the air-friction parameter, and *S*_d_ is the surface element on the disc. Integrating over the total surface area of the paper disc of width *w*, we obtain the following expression for the drag torque: 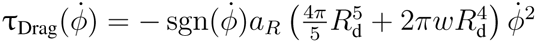.

The last twisting term (τ_Twist_) accounts for many forces in the paperfuge dynamics contributed through the resistance to twisting by the string. Since the strings have low stiffness in both bending and torsion, they undergo twist far beyond the zero-twist point (Fig. S6), to form supercoiled structures, as seen in Fig 1C (inset). To account for this twist resistance, we define an empirical equation, based on the following observations (inset Fig. 2A, Fig. S6, and movie S3): (i) at *ϕ* = 0, we don’t expect any resistive torque on the disc from the string, (ii) at *ϕ*_crit_, the string is at a geometric critical twist, and further compression is modeled as a linear spring force, and (iii) *ϕ* cannot exceed some maximum value *ϕ*_max_. Using these conditions, we define a function of the form, 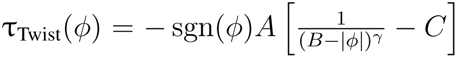, with a twisting parameter (*γ*) which we fit using experimental data (Fig. S6). *A, B, C* are found using the three constraints described above, resulting in the complete expression 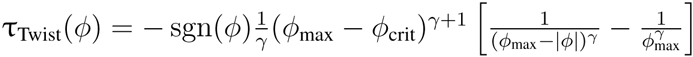.

Knowing the expression for the three torque terms, we numerically solve the differential equation to quantify the dynamics of the buzzer toy (Fig. 2A). Separating the individual contributions of the torque allow us to see the three stages where each torque term dominates in the cycle. τ_Input_ dominates during the unwinding phase, followed by τ_Drag_, which reaches its maximum value at 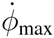. τ_Twist_ increases drastically at the end of the winding phase 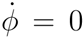, bringing the disc to a momentary halt, 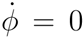.

By simultaneously measuring applied force and rotational velocity, we quantify the two unknown parameters in our model: drag coefficient (*a_R_*) and twisting parameter (*γ*), respectively. Using a high-speed camera (*f_rec_* = 5, 000 fps) synchronized with a force transducer, we obtain a time-series of the force actuating the paperfuge (Fig. 2B). By using a simple input function for the force (Fig. S7) in our model (Eq. 1, Fig. 2C i), we accurately predict the dynamics of the paperfuge (Fig. 2C ii-iii). We further utilize the predictive power of our model to develop a design landscape and compare with our experimental data (in red circles, Fig. 2D) and thus validate our model. We also highlight the dependence of 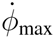 and *f*_0_ on different physical parameters, such as string radius and length, and the disc radius and width (Fig. S8, Tables S2 and S4).

With our validated model, we can now optimize the function of paperfuge for centrifugation. Centrifuges are characterized by the “relative centrifugal force” (RCF) they can generate, defined as 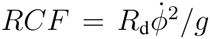 where *g* is acceleration due to gravity. The paperfuge is an oscillatory centrifuge, which changes directions in a periodic manner. Thus, we define an effective RCF as 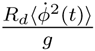 (Fig. S9) that yields an *RCF* ∼ 10, 000 g for a paperfuge with *R_d_* = 50 mm (see SI Fig. S9 for a table of RCF values for various paperfuge sizes).

Next, we demonstrate how the paperfuge can be utilized as a field-portable, ultra-low cost centrifugation tool (Fig. 3A,B, Fig. S2 and movie S5). We implement several safety measures to make our device usable in the field (see SI). We fill capillaries with 20 µL of whole human blood (from a finger-prick) and spin them on the paperfuge, revealing complete separation of plasma from red blood cells (RBC) within 1.5 mins (Fig. 3C, D). The volume fraction of RBCs provides a direct readout for hematocrit values (0.43, at *t* = 1.5 mins), which is a measure used for diagnosing anemia. The hematocrit value is in good agreement with control experiments conducted simultaneously on commercial electric-centrifuges (see SI). Furthermore, the resulting pure plasma can be easily retrieved for use with other RDTs.

**Figure 3.**
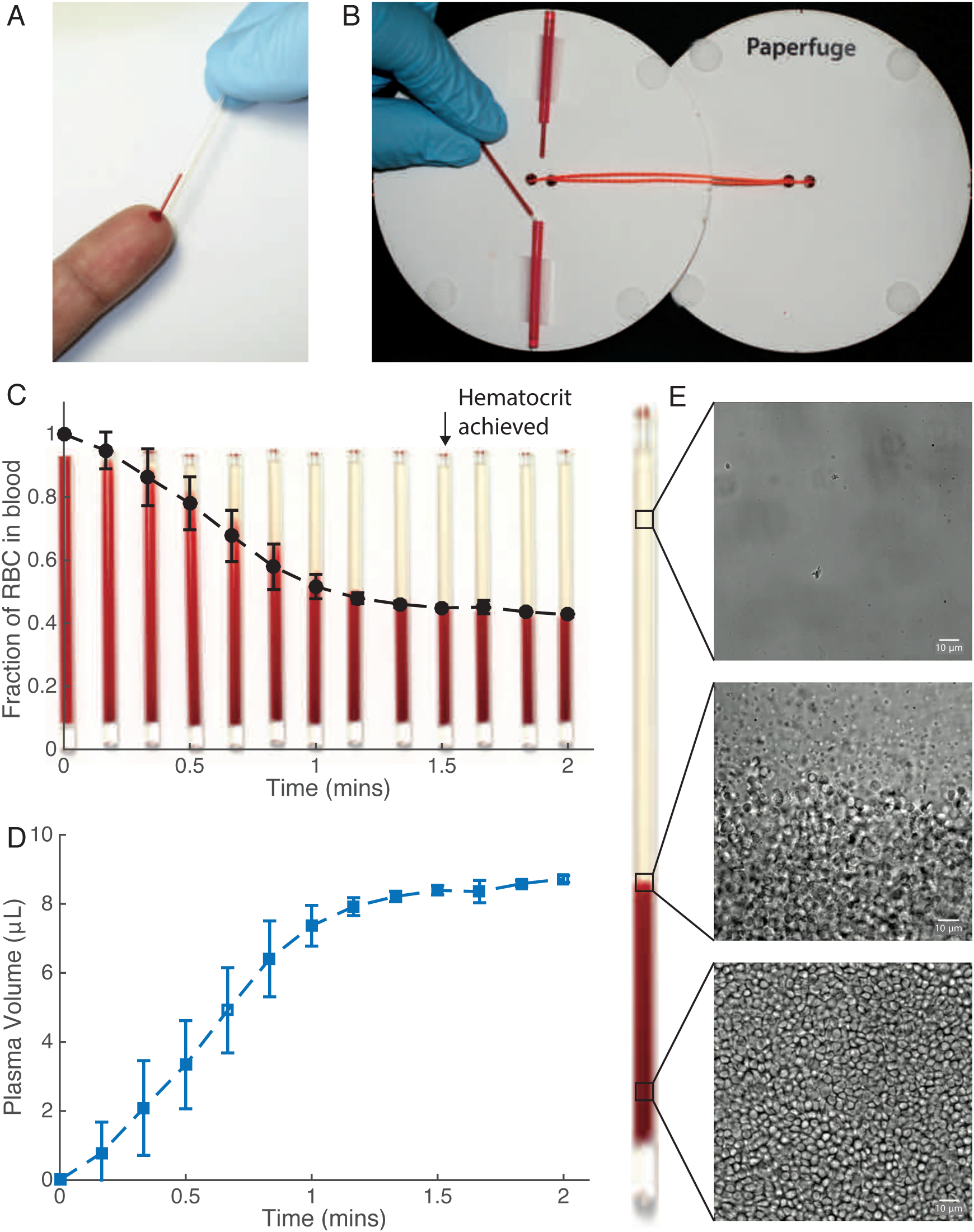
Paperfuge as a 20-cent diagnostic device for hematocrit analysis. The high rotational speeds of the paperfuge can be exploited in diagnostic sample preparation as an ultra-low cost centrifugation apparatus. **A-B)** 20 µL of human blood are loaded in plastic capillaries and placed into hollow plastic capillary holders (ends sealed) attached to paperfuge. **C)** Kinetics of RBC and plasma separation. The standard deviation on each data-point is calculated from over 8 separate trials conducted by two users (one male, one female) highlighting the consistency and reproducibility of the hematocrit values. **D)** We show that 8 µL of blood plasma (per capillary) can be separated from whole blood in less than 1.5 mins. **E)** The quality of the plasma is evaluated using microscopy and reveals 100% pure plasma.

Finally, we show the breadth of possibilities and applications of the paperfuge platform. Using a QBC capillary and float system, we show that 15 mins of spinning on a paperfuge can successfully separate the buffy coat (Fig. 4A). This expanded region can be easily used for identification of haematoparasites for infectious diseases such as malaria and African trypanosomiasis using a microscope (Fig. 4A, Table S1) (16). The simplicity and robustness of the paperfuge device makes it possible to design and construct devices from materials beyond paper, including wood, plastic and polymer. Using a desktop 3D printer (Form 2, Formlabs), we rapidly print light-weight (20 g) prototypes of different “3D-fuges” that spin at speeds of ∼ 10k rpm (Fig. 4B). These further open opportunities to mass-manufacture millions of centrifuges using injection moulding techniques. Moreover, we demonstrate centrifugal microfluidics in a “PDMS-fuge” (disc made from Polydimethylsiloxane (PDMS), Fig. 4C). This opens up possibilities to design integrated lab-on-a-chip devices that do not require external pumps or electricity (17). Since soft lithography requires fabrication infrastructure, we show that inexpensive plastic-tape microfluidics can be used with the paperfuge. We create two-dimensional plastic slides and demonstrate plasma isolation in 2 mins, which can be further imaged under a microscope without perturbing the sample (Fig. 4D).

**Figure 4.**
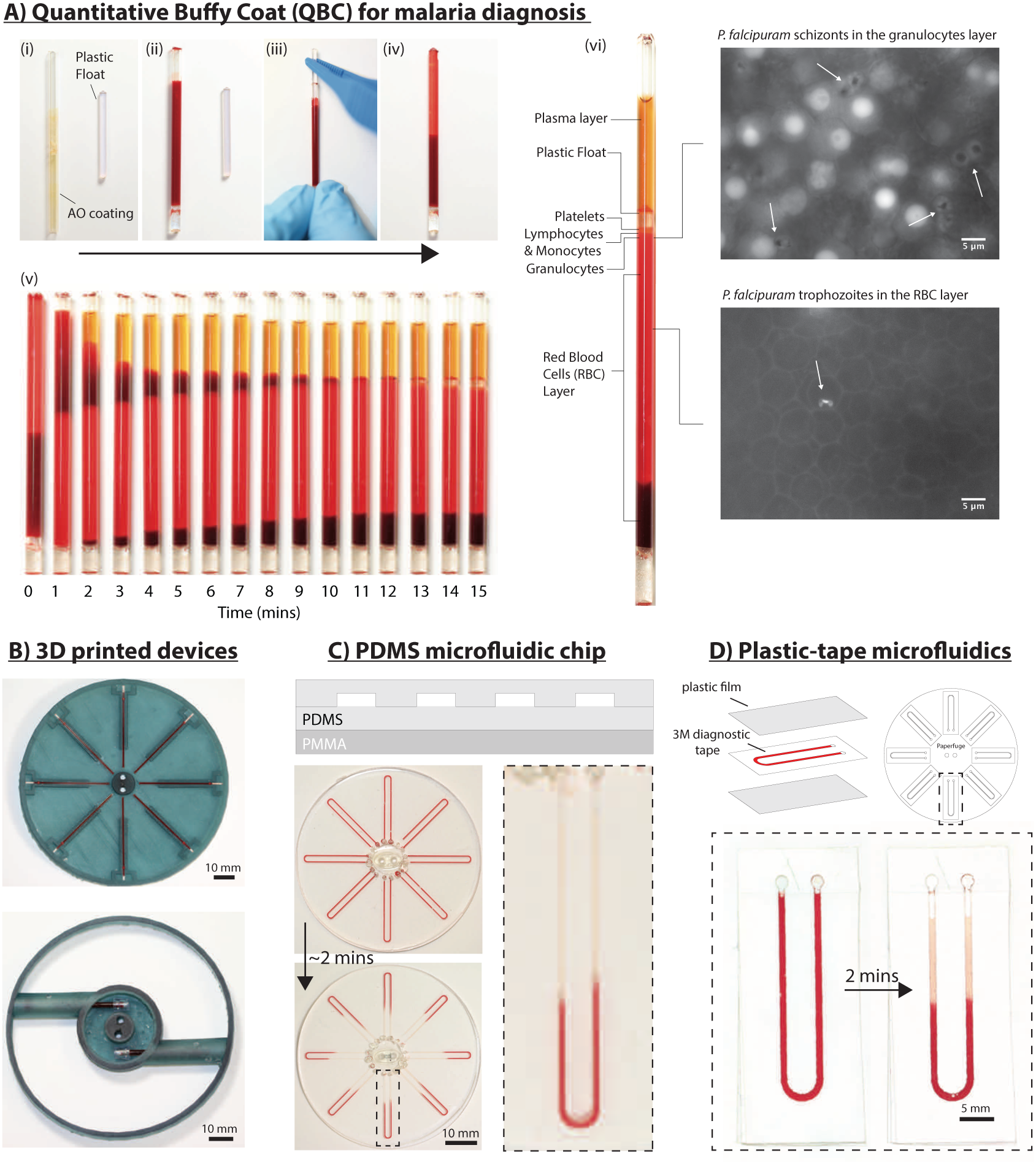
Paperfuge applications and design landscape. **A) Quantitative Buffy Coat (QBC) for malaria diagnosis)** (i-iv) Commercial QBC capillary (coated with Acrydine Orange dye) and solid cylindrical float can be utilized on the paperfuge to achieve density-based stratification of blood components. Snap-shots of separation are shown in (v): at *t* = 0, the float is inserted into the top of the capillary. As soon as spinning commences, the float settles to the bottom. Over a period of 15 mins, it rises to the plasma-RBC interface due to the difference in density, expanding the buffy-coat region. (vi) Using *Plasmodium falcipuram* infected blood, we can rapidly visualize the parasite using a fluorescent microscope in specific density-layers. **B) 3D printed devices** Light-weight polymeric materials can be rapidly printed for generating functional low-cost “3D-fuges”. The two prototypes shown achieve speeds of ∼ 10k rpm and weigh *<* 20 g. **C) PDMS centrifugal microfluidics** Using soft lithography, PDMS devices are fabricated to exploit centrifugal microfluidics. This “PDMSfuge” spins at 15k rpm and plasma is separated in 2 mins from 10µL of blood. **D) Plastic-tape microfluidics** Schematic showing two-dimensional “slides” constructed using 3M Diagnostic Tape that can easily be attached to a paperfuge. A 2-D slide with a u-shaped channel shows plasma separation from 15µL whole blood in 2 mins.

## Conclusion

The in-depth explorations of a simple toy here provide broad inspiration for developing human-powered, instrument-free POC devices. The variety of materials (paper, polymers) and rich design landscape offer potential for electricity-free integrated POC devices. The choice of paper as a substrate further opens opportunities for incorporating origami-based geometries, embedding optics (18), paper-based microfluidics (19) and ultimately integrated lateral flow rapid diagnostic as-says (20). Moreover, by exploiting the unique oscillatory dynamics of a paperfuge, new separation protocols can be explored that have theoretically been predicted but not confirmed empirically (21). The simplicity in manufacturing of our proposed solution will enable immediate mass distribution of a solution urgently needed in the field. Ultimately, our present work serves as an example of frugal science: leveraging the complex physics of a simple toy, for global health applications.

## Materials and Methods

### Paperfuge

The paperfuge is composed of two card-stock paper discs (SI Fig. 2A). Braided fishing line (Dyneema, MagicShield) is used for the strings to provide high tensile strength. Common wood/PVC pipe is used for the handles. We used drinking straws to create safe and easy mounting of the capillaries. The straws are sealed using epoxy (J-B Weld 50133, Plastic Bonder) to act as a secondary containment for accidental leakages from the capillary. First, two equal circle-shaped discs (*R_d_* = 50 mm) are printed on the paper using a laser printer (Epilog Mini Laser). In the center of each disc, there are two small circular holes (diameter = 3 mm), spaced 2.5 mm apart. During spinning motion, the strings cause extensive friction at the point of contact with the paper. To reduce wear, we use two 3 mm thick, acrylic ovals (major-axis = 13 mm, minor-axis = 6 mm) with two small circular holes (diameter = 3 mm), spaced 2.5 mm apart. These acrylic ovals are taped to the center of the outer face of each disc, such that the holes on the acrylic align with the holes on the disc. Two straws are cut down to 40 mm long pieces. One end of each straw is sealed with a drop of epoxy and allowed to dry for at least 20 mins at room temperature. The straws are glued at opposite edges of the inner face of one of the discs, with the epoxy-closed ends pointing outwards. The four velcro pieces are placed 90 degrees apart along the edges of the circular disc. These velcro pieces stick the two paper-discs, creating a streamlined surface, and thus reduce air-drag. The string is threaded through the holes in the center of the discs, and each end of the string is tied around the dowel. The two discs are then attached using velcro, covering the straws (SI Fig. 2B). The paperfuge is spun by holding one dowel in each hand (SI Fig. 2C).

### User Safety Measures

The paperfuge has three independent safety mechanisms to protect the user from accidental exposure to blood. First, the capillaries employed are made of plastic that are shatter-proof. Second, the capillaries are inserted into sealed-straw holders that can contain accidental leaks. Last, the paperfuge has two discs, one disc with the straw capillary holders and one disc as a cover, held together using velcro strips. This cover serves as an additional safety measure to prevent blood exposure to the user.

### Plasma Separation from Whole Human Blood

In all the experiments conducted in this study, human blood samples were purchased from ZenBIO Inc., and stored in the fridge until used. Plastic capillaries (40 mm long, SafeCrit heparinized plastic microhematocrit tube, 22274913, Fisher Scientific Inc.) are filled with blood, up to the marked line. One end of the capillary is sealed using epoxy (5-Min Epoxy) and allowed to sit for at least 20 mins at room temperature to completely seal the capillary. The capillary is then mounted into the straw, with the sealed end facing away from the center.

We have conducted more than 50 trials, with two operators (1 male, 1 female). The hematocrit results obtained in 1.5 mins on the paperfuge (PCV=0.43) are comparable to the results obtained using a commercial centrifuge (PCV=0.47) in 2 mins, made by Beckman Coulter (Critspin). The Critspin centrifuges the blood at a speed of 16,000 rpm (13,700 g) and costs $700. The paperfuge (with two loaded capillaries) spins at at a maximum speed of ∼ 20, 000 rpm (∼ 10, 000g) and costs 20 cents.

### Quantitative Buffy Coat (QBC) Protocol

For Quantitative Buffy Coat (QBC) analysis using the paperfuge, capillaries coated with Acrydine Orange dye, from QBC Malaria Test kit (Drucker Diagnostics) were used. The kit also contains a Precision Plastic Float that suspends in the buffy coat layer post-centrifugation, expanding it for further visualization via brightfield and fluorescent microscopy. To fit on a *R_d_* = 50 mm paperfuge, we cut the QBC capillary using a diamond cutter. Next, the capillary is filled with 30 µL of blood sample (spiked with 7.5% *P. falcipuram* parasitemia) and capillary sealed using epoxy at one end. The float is then fully inserted into the open end of the capillary using plastic forceps. The closed-end of the capillary is then inserted into the straw-holder and the paperfuge is then spun for 15 mins.

Over 10 trials of centrifugation were conducted using the QBC capillaries. After the first minute, the float immediately moves to the bottom of the capillary, and gradually rises due to its specific density, as the blood separates into plasma and RBCs within 15 mins. The buffy coat region is then examined using microscopy. Since we only use healthy human blood, no parasites are observed. However, QBC has been successfully shown to diagnose a large number of infectious diseases (see SI Table 1). Thus, we show proof-of-concept that QBC analysis can be conducted using the paperfuge for disease diagnostics.

### PDMSfuge

The PDMS-fuge is fabricated using the soft lithography technique. First, SU-8 50 (MicroChem Corp.) resist is spin-coated on a wafer, followed by pre-baking for 10 mins, resulting in a 110 µm thick layer. This is further baked for 30 mins to densify the resist film. The resulting resist is then exposed to UV-light for 18 s, which is post-baked for 10 mins to selectively cross-link the exposed parts. Finally, the unexposed material is washed away with the SU-8 developer, to obtain the master mold. To fabricate the channels, a PDMS mixture of 20 : 1 base and curing agent is poured onto the mold. A supporting PDMS/PMMA based structure is also made by pouring a 5: 1 mixture onto circular PMMA disc (1.5 mm thick). Finally, both layers are bonded by incubation at 65°C overnight. Each U-channel holds up to ∼ 10µL of liquid.

### Plastic-tape microfluidics

Plastic-tape microfluidics provide an easy to use, low-cost, disposal platform for centrifuging biological samples. The slides were assembled using double-sided tape (3M 9965 Double-Coated Polyester Diagnostic Tape) and thin plastic films (3M pp2500 Transparency Film). First, U-shaped channels were laser-printed onto two sheets of double-sided tape. Each side of the tape was then adhered to a thin plastic film, and the entire microfluidic channel was clamped for 15 mins to seal the sides of the channel. A drop of blood (∼ 10µL) was applied to the top of the channel, and capillary-action drew the blood into the channel in a few seconds. These slides were then mounted on a paperfuge and spun normally. Alternatively, paper can be used as the channel layer, and single-sided tape can be used to seal from both sides.

### High-speed Dynamics Setup

To measure the dynamic force, we used an S-type load cell (CZL301C, Phidgets Inc.). The load cell was connected to an Arduino (RedBoard, SparkFun Electronics) using a load-amplifying circuit board (HX711, SparkFun Elec). A custom code was written to calibrate the load-cell with known weights, and measure the force output when connected to one end of the paperfuge setup. High speed videos were recorded using two high-speed cameras. Analysis was done using custom script written in Matlab.

## Supplementary Material

- Supplementary Text
- Supplementary Theory and Equations
- Fig. S1 Antique buzzer toys
- Fig. S2 Paperfuge materials
- Fig. S3 High-speed setup
- Fig. S4 Schematic for theoretical model
- Fig. S5 Evaluating slip, no-slip and transition boundary conditions
- Fig. S6 Twisting torque
- Fig. S7 Theoretical input force
- Fig. S8 Theoretical design landscape for paperfuge scaling
- Fig. S9 Scaling RCF for paperfuge
- Table. S1 List of human infectious diseases diagnosable using QBC technique
- Table. S2 Parameter values in scaling analysis
- Table. S3 Experimental parameters for various paperfuge sizes
- Table. S4 List of parameters in theoretical models
- Movie S1 High speed dynamics of paperfuge
- Movie S2 Measuring rotational speed
- Movie S3 Twisting torque condition
- Movie S4 Comparison of experiment and theory
- Movie S5 Separation of plasma from whole blood using paperfuge

## Acknowledgments

We thank all members of the PrakashLab for useful feedback, Felix Hol for assistance with soft lithography fabrication, and Katrina Hong (Yeh Lab, Stanford University) for assistance with malaria samples. MSB acknowledges fellowship support from Stanford School of Medicine Dean’s Postdoctoral Fellowship and SPADA. MP acknowledges support from Pew Foundation, Moore Foundation, SPADA and NIH New Innovators Award. This work was supported by the Stanford Clinical and Translational Science Award (CTSA) to Spectrum (UL1 TR001085). The CTSA program is led by the National Center for Advancing Translational Sciences (NCATS) at the National Institutes of Health (NIH). The content is solely the responsibility of the authors and does not necessarily represent the official views of the NIH.

## Author contributions

M.S.B and M.P. designed research; M.S.B, B.B.,C.C., G.K. and A.J. performed and analyzed the research; all authors wrote the paper.

## Competing Interests

The authors declare they have no competing interests.

